# Splicing inactivation generates hybrid mRNA-snoRNA transcripts targeted by cytoplasmic RNA decay

**DOI:** 10.1101/2022.02.02.478905

**Authors:** Yanru Liu, Samuel DeMario, Kevin He, Michelle R. Gibbs, Keaton W. Barr, Guillaume F. Chanfreau

**Affiliations:** Department of Chemistry and Biochemistry, UCLA, Los Angeles, CA 90095, United States of America; Department of Molecular Biology and Genetics, Cornell University, Ithaca, NY 14853-2703; Molecular Biology Institute, UCLA, Los Angeles, CA 90095, United States of America

**Keywords:** snoRNA, splicing, introns, exosome, exonuclease

## Abstract

Many small nucleolar RNAs (snoRNA)s are processed from introns of host genes, but the importance of splicing for proper biogenesis and the fate of the snoRNAs is not well understood. Here we show that inactivation of splicing factors or mutation of splicing signals leads to the accumulation of partially processed hybrid mRNA-snoRNA transcripts (hmsnoRNA). HmsnoRNAs are processed to the mature 3′-ends of the snoRNAs by the nuclear exosome and bound by snoRNP proteins. HmsnoRNAs are unaffected by translation-coupled RNA quality control pathways, but they are degraded by the major cytoplasmic exonuclease Xrn1p due to their mRNA-like 5′-extensions. These results show that completion of splicing is required to promote complete and accurate processing of intron-encoded snoRNAs and that splicing defects lead to degradation of hybrid mRNA-snoRNA species by cytoplasmic decay, underscoring the importance of splicing for the biogenesis of intron encoded snoRNAs.

**Significance Statement:** Small nucleolar RNAs mediate modifications of nucleosides within ribosomal RNAs, which are necessary for proper ribosomal function and translation. Many snoRNAs are encoded within introns of host genes and accurate biogenesis of these small RNAs is required to produce functional snoRNAs. The work presented here shows that when the splicing reactions are inactivated, snoRNAs undergo a distinct biogenesis pathway which leads to the production of aberrant hybrid RNAs that contain both mRNAs and small RNAs components of the host genes. While snoRNAs are primarily found in the nucleolus, these hybrid RNAs are degraded by the cytoplasmic mRNA degradation pathway. These results demonstrate the importance of splicing to promote accurate snoRNA processing and prevent the production of aberrant mRNA-snoRNA hybrids.

## Introduction

Small nucleolar RNAs (snoRNAs) are non-coding RNAs that guide snoRNP-mediated 2′-O-methylation or pseudouridylation of pre-ribosomal RNAs precursors and other stable RNAs (1),2,3. These snoRNAs are classified in two major families: the C/D and H/ACA classes, which guide 2′-O-methylation and pseudouridylation of their substrates, respectively. The importance of correct snoRNA expression is underscored by the fact that defects in snoRNA metabolism are linked to multiple pathologic processes including cancer (4, 5), Prader-Willi syndrome (6), and metabolic stress (7). SnoRNAs are found in many different genomic contexts (2, 8). In mammalian genomes, many snoRNAs are present in the introns of host genes, but they can also be generated from lncRNAs. In plants, snoRNAs are often generated from polycistronic transcription units(8). In the budding yeast *S*.*cerevisiae* which has been used extensively to study the mechanisms of snoRNA biogenesis and processing, snoRNA are expressed either from independently transcribed genes, polycistronic snoRNA precursors, or from introns (2, 8). The synthesis of mature snoRNAs from intronic sequences is thought to occur primarily through splicing of the host gene pre-mRNA. *In vitro* and *in vivo* studies have shown that splicing results in accurate processing of intron-encoded snoRNA in mammalian cells (9). The current model is that intron-encoded snoRNAs are generated by exonucleolytic trimming of the excised linear introns after completion of the splicing reaction and debranching of the lariat introns. In support of this model, inactivation of the *S*.*cerevisiae* debranching enzyme Dbr1p results in the accumulation of lariat intron species containing the snoRNAs (10), showing that debranching of the excised intron is critical for processing. However, the production of mature snoRNA can still occur inefficiently in the absence of the debranching enzyme(10), because random hydrolytic cleavage of the lariat intron exposes the cleaved intron to processing by exonucleases or by cleavage by the RNase III enzyme Rnt1p(11). In addition, some studies have reported splicing-independent processing of intron-encoded snoRNAs (12), either by endonucleolytic cleavage of the precursors (11, 13), or by exonucleolytic processing (14). In *S*.*cerevisiae*, the RNase P endonuclease has also been proposed to initiate a processing pathway independent from splicing, by cleaving unspliced pre-mRNAs that host box C/D snoRNAs (15). Finally snoRNAs can be found associated with stable lariats(16) and processing of introns containing multiple snoRNAs can generate sno-lncRNAs, which correspond to partially processed introns that contain two snoRNAs linked by an intronic segment (17).

Despite the known importance of splicing for the processing of intron-encoded snoRNAs, it is unclear how defects in the spliceosome machinery may impact the fate of intron-encoded snoRNAs. This is an important question, as recent work has shown that defects in 5′-end processing of independently transcribed snoRNAs can result in mislocalization of the unprocessed snoRNAs in budding yeast(18). Strikingly, very little is known about the impact of spliceosome defects on snoRNA expression. Haploinsufficiency of the core snRNP protein SmD3 results in a reduction in the levels of intron-encoded snoRNAs(19), but the specific molecular effects of this mutation on the biogenesis pathway of intron-encoded snoRNAs have not been investigated. In this study, we have analyzed the impact of inactivating splicing using *trans* and *cis*-acting splicing mutants on the production of intron-encoded snoRNAs in the yeast *S*.*cerevisiae*. We show that inactivating splicing factors involved in different steps of the spliceosome cycle, or mutating the splicing signal of a host gene result in the accumulation of aberrant, hybrid mRNA-snoRNAs species which share some of the hallmarks of mature snoRNAs, but are degraded by the cytoplasmic decay. These results highlight the importance of splicing for determining the fate of intron-encoded snoRNAs and show that incorrectly processed intron-encoded snoRNAs are degraded by the general mRNA decay pathway. Our results may also provide some insights onto possible additional effects of genetic diseases that affect splicing factors.

## Results

### Inactivation of splicing factors at different stage of the spliceosome cycle results in the accumulation of hybrid mRNA-snoRNA forms of *NOG2/snR191*

To investigate the importance of the splicing reaction for the processing of intron-encoded snoRNAs, we used the anchor away (AA) technique(20), which promotes the rapid export of endogenously FRB (*FKBP12*-rapamycin binding domain)-tagged splicing factors to the cytoplasm after addition of rapamycin (20). This results in the nuclear depletion of these factors and in the inactivation of their nuclear functions *in vivo*. The genetic background used for these experiments alleviates the toxic effects of rapamycin and its downstream effects on gene regulation (20). To analyze the contribution of splicing factors involved at different steps of the spliceosome cycle, we used strains expressing FRB-tagged versions of Prp5p, Prp28p, Prp16p, Prp18p, Slu7p, and Prp22p. These proteins are involved in various steps of splicing (21), from spliceosome assembly (Prp5p, Prp28p) to the second catalytic step (Prp16p, Prp18p, Slu7p, Prp22p) and spliceosome recycling (Prp22p). We first analyzed expression of the *NOG2* gene which contains the H/ACA snoRNA snR191 in its intron (22) by Northern blot (Figure 1A). After nuclear depletion of each of these splicing factors, a probe hybridizing to the exon1 of *NOG2* detected the unspliced precursors, but also RNAs migrating faster than the spliced mRNAs (labeled hms on Figure 1B). Hms RNAs were detected in all the anchor away strains treated with rapamycin, but not prior to rapamycin treatment. We first hypothesized that these species might correspond to cleaved 5′-exons; however, based on their estimated size (∼1100nt), these species are too large to correspond to free 5′ exons. In addition, these RNAs were also detected after anchoring away proteins involved prior to the first catalytic step (eg. Prp5p, Prp28p). Furthermore, species migrating faster than the hms RNAs and whose size matches those of cleaved 5′exons (∼850nt) were detected only in strains in which second step splicing factors are anchored away (Slu7p, Prp16p, Prp18p, Prp22p), but not in strains in which splicing factors involved prior to the first step are depleted from the nucleus (Prp5p; Prp28p). These fastest migrating species correspond to cleaved 5′-exons that fail to undergo splicing because of second step defects and are labeled E1 in all figures.

**Figure 1.**
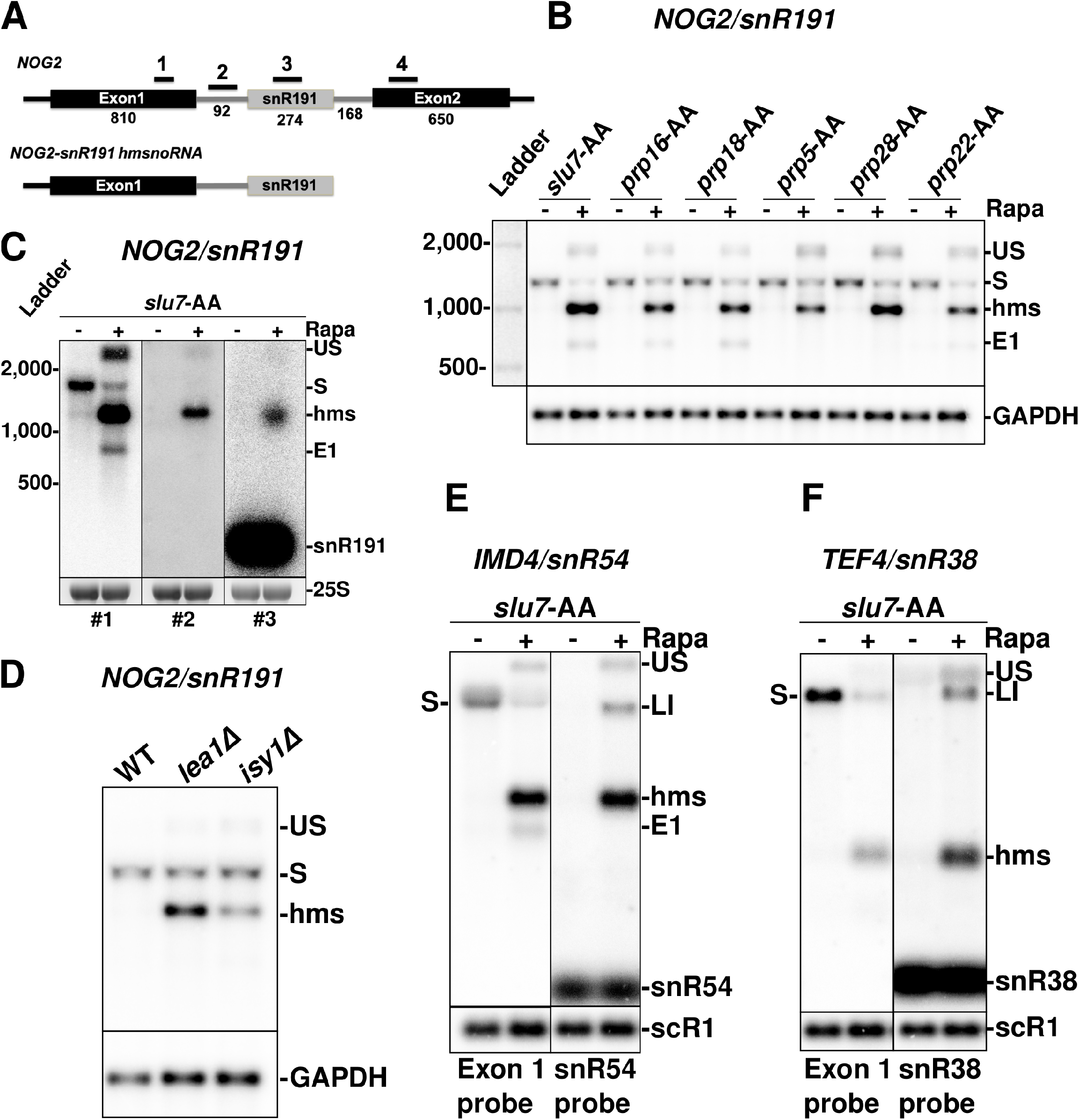
Hybrid forms of intron-encoded snoRNAs are produced upon splicing inactivation. **A.** Schematic structure of the *NOG2* gene encoding the *snR191* H/ACA snoRNA in its intron, and proposed structure of the *NOG2-snR191* hmsnoRNA. Exons are represented by black boxes and the mature snoRNA by a gray box. Intronic sequences are shown as gray lines. Numbers indicate the length in nucleotides of the different exonic and intronic regions and of the snoRNA. UTR lengths are not included. Black lines with numbers 1-3 indicate the approximate locations of the different riboprobes used for this figure. Boxes and line lengths are not to scale. **B.** Northern blot analysis of *NOG2/snR191* in strains expressing anchor-away (AA) FRB-tagged versions of the Slu7p, Prp16p, Prp18p, Prp5p, Prp28p and Prp22p splicing factors. Each FRB tagged strain was grown in normal medium, and then spun down and resuspended in either fresh normal media or shifted to media containing rapamycin for 1hr to promote export of the tagged splicing factor out of the nucleus. Probe#1 was used for the top panel. GAPDH was used as a loading control. US = unspliced pre-mRNA; S = Spliced mRNA; hms = hmsnoRNA. E1 = cleaved exon1. **C.** Mapping of the hmsnoRNA forms of *snR191* using different riboprobes. Shown are northern blots of *NOG2/snR191* using riboprobes 1,2 and 3 shown in (A) of RNAs extracted from the Slu7-AA strain before or after 1hr treatment with rapamycin. An ethidium bromide staining of the 25S rRNA is shown as a loading control. **D.** WT Northern blot analysis of *NOG2* in strains in WT and *lea1*Δ or *isy1*Δ deletion strains. Legend as in (B). **E.** Northern blot analysis of *IMD4/snR54* in RNAs extracted from the Slu7p-AA strain before or after treatment with rapamycin for one hour. Shown are northern blots using probes hybridizing to the 5′-exon or to the mature snoRNA. Labeling of the different species as in (B). *scR1* was used as a loading control. **F.** Northern blot analysis of *TEF4/snR38* in the Slu7p-AA strain Legends as in (B). LI = Lariat Intron-Exon2 intermediate.

Further mapping of the hms RNAs using probes hybridizing to the different regions of *NOG2-snR191* gene (Figure 1A) showed that these species also hybridize to probes complementary to the mature *snR191* snoRNA and to the intronic region preceding the snoRNA (Figure 1C). Based on their approximate size (∼1100 nucleotides) and their hybridization patterns, we hypothesized that these RNAs correspond to hybrid mRNA-snoRNA (hmsnoRNA) containing the mRNA 5′-exon, the intronic segment upstream of the snoRNA, and the entire snoRNA sequence (schematic representation in Figure 1A). The combination of estimated sizes and probe hybridization patterns was consistent with a 3′-end close to that of the mature snoRNA 3′-end. To test this idea, we mapped the 3′-end of the *NOG2-snR191* hmsnoRNA using a modified 3′-RACE protocol after i*n vitro* polyadenylation of total RNAs. Sequencing of the 3′-RACE products showed that the *snR191* hmsnoRNA 3′-ends match precisely those of mature snoRNAs (Figure S1). RNAs similar in size to the hms species also accumulated in mutants carrying deletions of the non-essential genes encoding the U2 snRNP component Lea1p(23) or the splicing fidelity factor Isy1p(24) (Figure 1D). Therefore, the production of hmsnoRNAs is caused by general splicing defects rather than by indirect effects of the anchor away process.

### HmsnoRNAs can be detected for most intron-encoded *S*.*cerevisiae* snoRNAs by northern blot and Nanopore long-read Sequencing

To extend the results described above for other snoRNAs, we analyzed RNAs accumulating in the Slu7p-AA strain prior to or after rapamycin treatment for several genes expressing box C/D snoRNAs from their introns: *IMD4*/*snR54* (Figure 1E), *TEF4*/*snR38* (Figure 1F) and *ASC1*/snR24 (Figure S2). For *IMD4*/*snR54* and *TEF4*/*snR38*, we detected the accumulation of hmsnoRNAs hybridizing to both exon1 and snoRNA probes (Figures 1E and 1F). Mapping of the *ASC1/snR24* hmsnoRNA using four probes hybridizing to different regions of *ASC1* (Figure S2A, S2B) showed that these species contain the first exon of *ASC1*, the first part of the intron and the entire snoRNA sequence that ends close to the mature 3′-end (Figure S2A,S2B). The estimated size and hybridization patterns of the *IMD4/snR54* and *TEF4/snR38* hmsnoRNAs (Figures 1E and 1F) suggest a similar architecture. Mapping of the 3′-end of the *IMD4/snR54* hmsnoRNA using the modified 3′-RACE protocol showed that the *IMD4/snR54* hmsnoRNA 3′-end is identical to that of the mature *snR54* (Figure S1). Thus, splicing inactivation results in the general accumulation of hybrid mRNA-snoRNA species for all intron-encoded snoRNA tested, which belong to both H/ACA and C/D families.

To perform a more comprehensive mapping of hmsnoRNA genome-wide, we performed a direct RNA long-read sequencing experiment on RNAs extracted from a Slu7-AA strain after addition of Rapamycin using the Oxford Nanopore platform. Prior analysis revealed that the majority of the *ASC1/snR24* hmsnoRNAs are not polyadenylated (Figure S2C), so the standard protocol to perform direct poly(A)+ RNA sequencing could not be used. We developed a protocol that used in vitro polyadenylation after rRNA depletion, which allowed detection of the hmsnoRNA by Nanopore sequencing. In the strain in which Slu7p was anchored away, hmsnoRNAs could be detected for *EFB1-snR18, ASC1-snR24, TEF4-snR38, IMD4-snR54, RPL7A-snR39* and *NOG2-snR191* (Figure 2), and *RPL7B-snR59* (Figure S3A). The architecture of hmsnoRNAs revealed by long read sequencing matched perfectly the predictions made above using northern blot, and confirmed that these species contain the 5′-exon, the beginning of the intron and the snoRNA up the mature 3′-ends. We also detected unspliced precursors and cleaved 5′-exons that accumulate due to the second step splicing defect (Figure 2). We did not detect any hmsnoRNA for the *RPS22B-snR44* gene (Figure S3B), likely because the presence of an RNase III cleavage site in the first intron of the unspliced precursors, which precludes the accumulation of Exon1-intron1-Exon2-snR44 species when unspliced species are generated (25). Our sequencing approach produced many reads for the box H/ACA *snR191* (Figure 2F) but failed to detect most mature box C/D snoRNA, with the exception of a few reads for *snR18* (Fig.2A).This lack of detection may be linked to the small size of box C/D snoRNAs which makes their detection challenging by Oxford Nanopore sequencing. Overall, these results show that after splicing inactivation, hmsnoRNA are generated from most intron-encoded snoRNAs in *S*.*cerevisiae* and the reads obtained confirm the general architecture of these RNAs.

**Figure 2.**
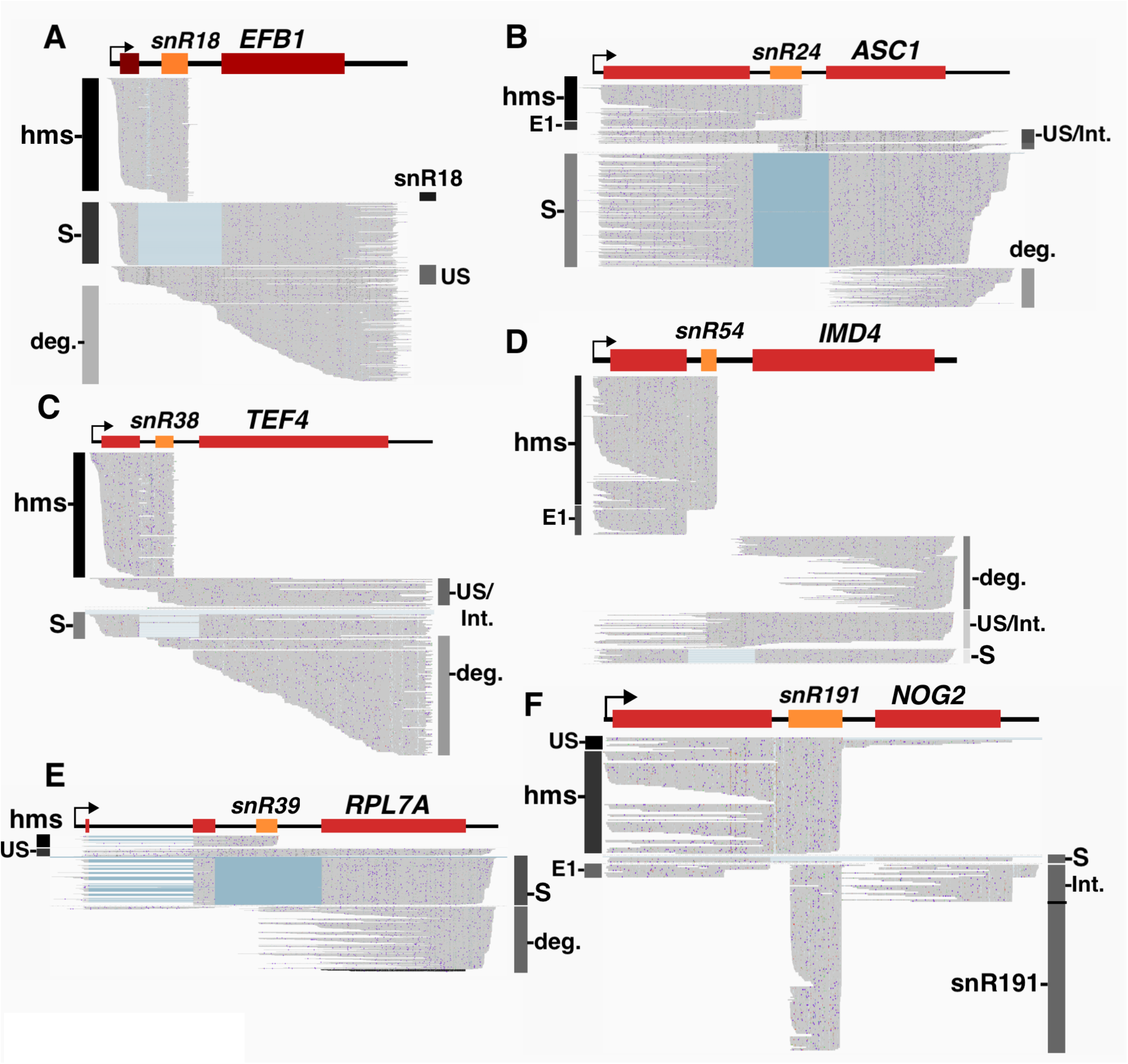
Detection of HmsnoRNAs by direct single molecule Oxford Nanopore RNA sequencing. For all the panels below, each sequencing read is represented by a horizontal gray line. RNA species are labeled as: hms = hmsnoRNA; US = unspliced; S = spliced mRNA; E1 = cleaved 5’exon; D= degradation intermediates; I = snoRNA processing intermediates. **A**. Oxford Nanopore sequencing reads obtained from a *slu7*-FRB tagged strain treated for one hour with Rapamycin for the *EFB1-snR18* locus. **B**. Same as (A) for *ASC1-snR24*. **C**. Same as (A) for *TEF4-snR38*. **D**. Same as (A) for *IMD4-snR54*. **E**. Same as (A) for *RPL7A-snR39*. **F**. Same as (A) for *NOG2-snR191*.

### HmsnoRNAs are processed at their 3′-ends by exonucleolytic trimming by the exosome

The previous results using 3′-RACE and Nanopore sequencing showed that the 3′-ends of hmsnoRNAs are identical to those of mature snoRNAs, which are generated by exonucleolytic trimming by the nuclear exosome (26). To test the hypothesis that hmsnoRNA acquire their 3′-end through exosome-mediated processing, we used an anchor away strain where the nuclear exosome component Rrp6p is FRB-tagged. Anchoring away Rrp6p provides a more effective method of inactivating the nuclear exosome than deleting the gene encoding Rrp6p (27), possibly because exporting Rrp6p to the cytoplasm may also result in the nuclear depletion of other nuclear exosome components. We then generated a strain in which Rrp6p and Prp18p or Slu7p could be co-anchored away simultaneously. Nuclear depletion of Rrp6p resulted in an increased accumulation of both the unspliced and spliced forms of *NOG2* (Figure 3A). This observation is consistent with previous studies that showed that the nuclear exosome actively degrades unspliced precursors and may limit the production of mature species(28–30). Hybridization with a probe complementary to snR191 detected the accumulation of shortly extended forms of snR191 in the Rrp6p-AA strain (Figure S4), consistent with the known role of the exosome in trimming intronic snoRNA mature 3′-ends (26).

**Figure 3.**
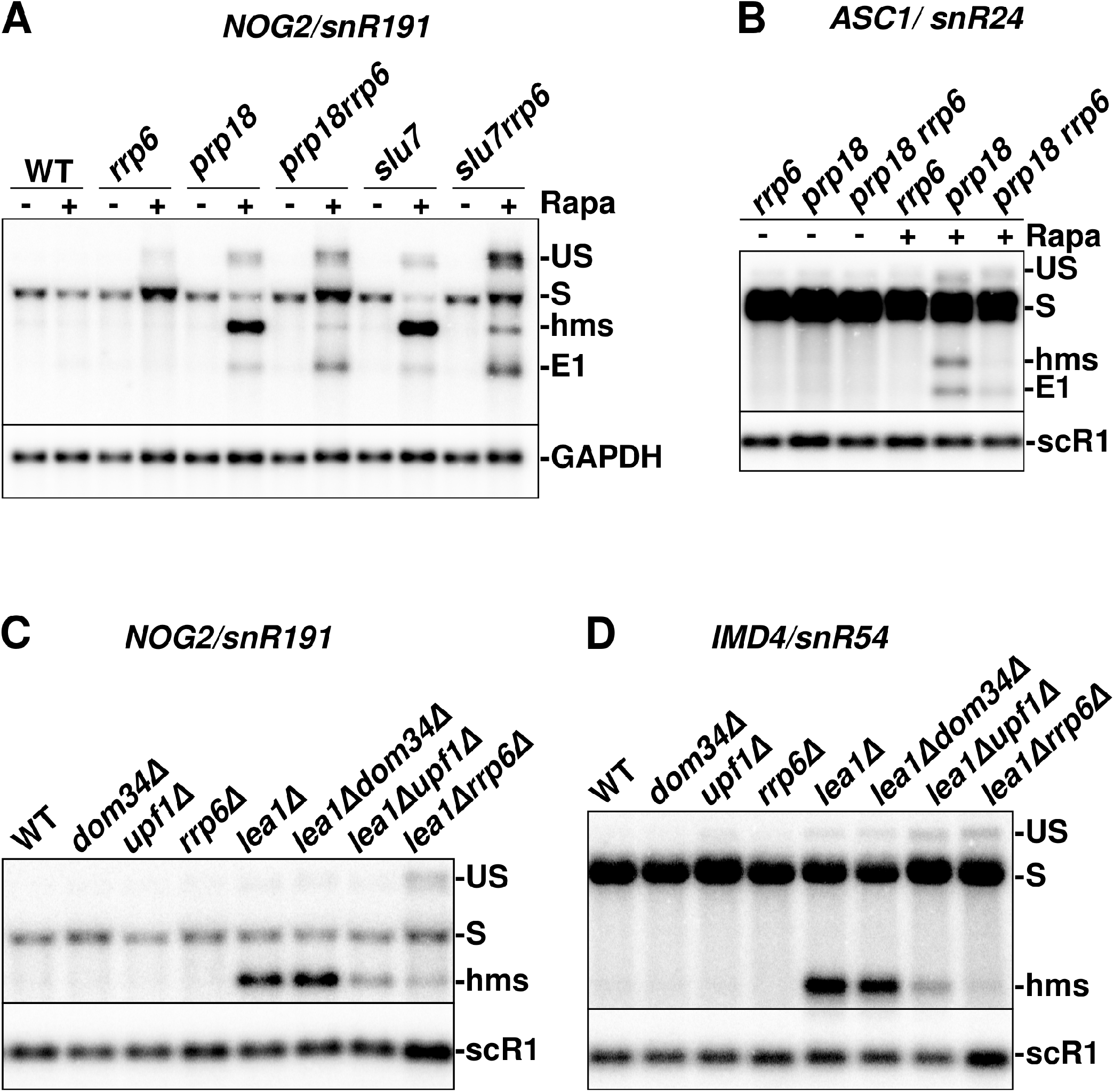
HmsnoRNAs acquire their 3′-ends through processing by the nuclear exosome but are mostly unaffected by cytoplasmic quality control pathways. **A**. Northern blot analysis of *NOG2/snR191* in strains expressing FRB-tagged, anchor-away versions of Rrp6p, Prp18p, Slu7p, or double anchor away versions. Strains in the name of the gene italicized indicate that the corresponding protein was FRB-tagged (eg *rrp6* = *rrp6-FRB*). The membrane was hybridized with a probe complementary to the exon1 of *NOG2*. GAPDH was used as a loading control. Legends as in Figure 1. **B**. Northern blot analysis of *ASC1/snR191* in strains expressing FRB-tagged, anchor-away versions of Rrp6p, Prp18p, or both Prp18p and Rrp6p. The probe used was complementary to the exon1 of *ASC1*. Scr1 was used as a loading control. Labeling of the species as in (A). **C**. Northern blot analysis of *NOG2/snR191* in strains carrying viable deletions of the genes encoding the splicing factor Lea1p, and/or the RNA degradation or processing factors Dom34p, Upf1p or Rrp6p. The probe used is complementary to the exon1 of NOG2. scR1 was used as a loading control. Labeling of the species as in (A). **D**. Analysis of IMD4/snR54 as described for NOG2/snR191 in panel C.

As shown previously, anchoring away Slu7p (Figure 3A and Figure S4) or Prp18p (Figure 3A), resulted in the accumulation of *NOG2-snR191* hmsnoRNAs. Strikingly, when Rrp6p was anchored away simultaneously with Prp18p or Slu7p, the accumulation of the hmsnoRNA decreased compared to what was observed when anchoring away these splicing factors alone (Figure 3A; see also Figure S4 for Slu7p). We observed a concomitant increase in the level of unspliced precursors (Figure 3A, Figure S4). This result suggested a precursor-to-product relationship between unspliced pre-mRNAs and hmsnoRNAs, and that hmsnoRNAs are generated by exonucleolytic trimming of unspliced precursors by the nuclear exosome.

To further demonstrate that hmsnoRNAs are processed to or near the 3′-ends of the mature snoRNAs by the exosome, we used strains where the gene encoding the non-essential U2 snRNP component Lea1p is deleted, along with a deletion of the gene encoding Rrp6p. Consistent with the results obtained above, the *rrp6*Δ knockout resulted in a reduction in the accumulation of *NOG2-snR191* hmsnoRNA species when coupled to the *lea1*Δ mutation, accompanied by an accumulation of unspliced precursors (Figure 3C). Overall, these results combined with the analyses described above show that hmsnoRNA are processed by the nuclear exosome to generate 3′-ends identical to those of the corresponding mature snoRNAs.

### HmsnoRNAs are not targeted by translation-dependent quality control pathways

Because hmsnoRNAs contain 5′-UTR and exon1 sequences, they exhibit mRNA-like features at their 5′-ends which suggest that they might be translated. However, they lack a poly(A) tail, which may limit translation efficiency. If hmsnoRNAs are translated, translation would stop shortly after the end of the 5′-exon sequence because of the occurrence of premature stop codons in the intronic sequences, which may trigger nonsense-mediated decay. NMD could potentially affect hmsnoRNAs, since NMD can occur in the absence of a poly(A) tail in yeast (31). Alternatively, the presence of secondary structures in the snoRNAs or the binding by snoRNP proteins (see below) may block ribosome progression and trigger No-Go Decay (NGD)(32). To investigate a potential targeting of hmsnoRNAs by these two RNA surveillance pathways, we knocked out the genes encoding the NMD factor Upf1p or the NGD factor Dom34p in the *lea1*Δ mutant and analyzed *NOG2-snR191* and *IMD4-snR54* by northern blot (Figure 3C, 3D). Eliminating Upf1p or Dom34p did not increase the levels of hmsnoRNAs in the *lea1*Δ mutant (Figures 3C and 3D). Instead, the *upf1*Δ knockout resulted in a decrease of the level of hmsoRNAs, while eliminating Dom34p showed no effect. As opposed to what was observed in the *lea1*Δ*rrp6*Δ mutant, the decreased accumulation of the *NOG2-snR191* hmsnoRNA detected in the *lea1*Δ*upf1*Δ mutant did not correlate with an increased accumulation of the unspliced precursors for snR191(Figure 3C), suggesting that Upf1p is not involved in the conversion of unspliced precursors into hmsnoRNAs. We detected some unspliced precursors accumulation for *IMD4* in the *lea1*Δ*upf1*Δ mutant (Figure 3D), but this was also detected in the *upf1*Δ mutant. While we do not fully understand the molecular basis for the decreased accumulation of hmsnoRNAs in the *lea1*Δ*upf1*Δ mutant, experiments described below using splicing signals mutations suggest that this might be the result of indirect effects. Overall, we conclude from these experiments that hmsnoRNAs are not targeted by translation-mediated RNA quality control pathways.

### HmsnoRNAs can be generated by a 5′ splice site mutation and are degraded by the cytoplasmic decay pathway involving Xrn1p

The previous results showed that inactivation of splicing factors resulted in the accumulation of hybrid mRNA-snoRNA species. However, we could not rule out that anchoring away splicing factors or deleting genes encoding non-essential splicing factors may result in indirect effects that may hamper the interpretation of our results. To alleviate these concerns, we used a plasmid-based system to introduce splice site mutations that would inhibit splicing and circumvent any general splicing defects. We generated a construct expressing the *NOG2-snR191* gene from a centromeric plasmid, which expressed *NOG2* at levels that exceeded the RNAs expressed from the endogenous *NOG2* gene (Figure S5). We did not detect any hmsnoRNA or any other aberrant species from the plasmid-borne wild-type version of *NOG2* showing that it is fully and accurately processed (Figure S5). We then generated two mutants designed to inactivate splicing of the plasmid borne *NOG2* transcripts: M1 is a mutation of the 5′-splice site (SS), and M2 is a double mutation that combines M1 with a branchpoint (BP) mutation (Figure 4A). Since the plasmid-borne versions are expressed at higher levels than the endogenous *NOG2* transcripts, and because *NOG2* is an essential gene, we used these plasmids in strains that also expressed the endogenous *NOG2* gene, which facilitated the genetic analysis. Transcripts expressed from the M1 and M2 plasmids accumulated in wild-type strains mostly as hmsnoRNAs, and to a lesser extent unspliced transcripts (Figure 4B). The temperature used to grow strains had some influence on the level of accumulation of hmsnoRNAs derived from these mutants, as hmsnoRNAs were more abundant when cells were grown at 20°C compared to 30°C (Figure S6).

**Figure 4.**
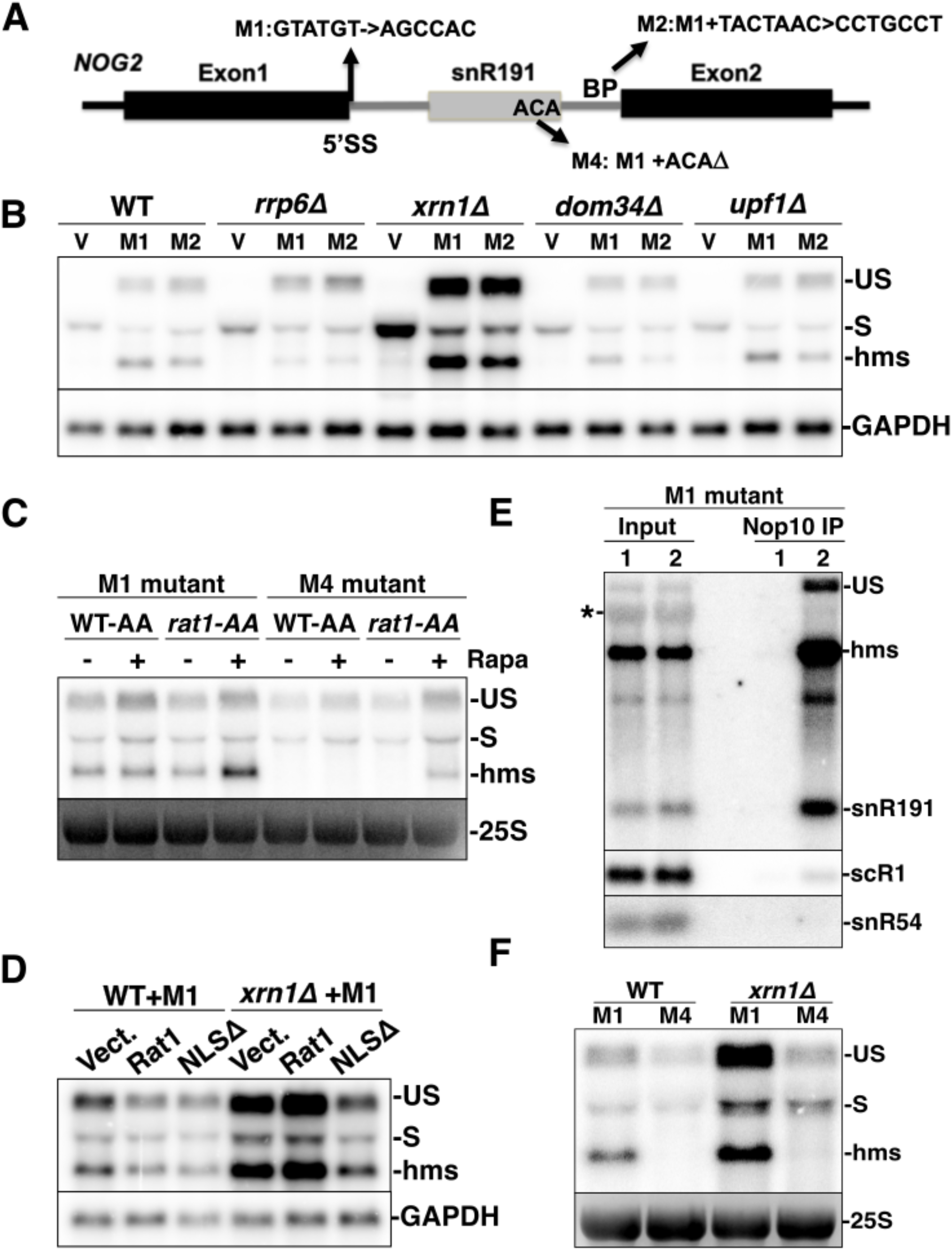
HmsnoRNAs generated by splice site mutations are degraded by Xrn1p and Rat1p and share some of the features of mature snoRNPs. **A**. Mutations introduced in the plasmid expressing *NOG2-snR191*. 5′SS = 5′ splice site; BP = branchpoint. **B**. *NOG2-snR191* hmsnoRNAs generated by splicing signal mutations are degraded by Xrn1p but unaffected by cytoplasmic quality control pathways. Wild-type (WT) or the indicated deletion mutants were transformed with the pUG35 vector (V) or the pUG35 plasmids expressing mutants M1 or M2 and RNAs extracted from these stains were analyzed by northern blot using a probe hybridizing to the exon1 of *NOG2*. GAPDH was used as a loading control. RNA species labeling as in Figure 1B. **C**. Impact of Rat1p inactivation on hmsnoRNA levels expressed from the M1 and M4 plasmids. A wild-type strain or a Rat1-FRB anchor away strain transformed with the M1or M4 expression plasmids were grown in medium in the absence or presence of Rapamycin (Rapa) for one hour and analyzed by northern blot as described in panel (B). 25S rRNA was used as a loading control. **D**. Impact of Rat1p delocalization on hmsnoRNA levels expressed from the M1 plasmid. A wild-type strain or an *xrn1Δ* mutant strain transformed with the M1 expression plasmid were transformed with an empty vector (vect.), a plasmid expressing wild-type Rat1p (Rat1) or a version of Rat1p with a mutated nuclear localization signal (NLSΔ). RNAs were analyzed by Northern blot as described in panel (B). **E**. NOG2-snR191 hmsnoRNAs generated by the M1 mutation are bound by the H/ACA snoRNP Nop10p. Shown are northern blots of RNAs extracted from cell extracts (Input) prepared from untagged wild-strain (Lane1) or ZZ-tagged Nop10 strain (Lane2) and of RNAs extracted after incubation of whole cell extracts from the same strains with IgG beads and washing (see methods). Membranes corresponding to independent purification experiments were hybridized to a probe complementary to the exon1 of NOG2 and to snR191 (top panel) or to probes hybridizing to scR1 or snR54 (bottom panels). The asterisk sign near the input lane indicates cross-hybridization of the probe to a ribosomal RNA. **F**. Destabilization of hmsnoRNAs caused by the deletion of the ACA box of plasmid-borne *NOG2-snR191*s cannot be rescued by inactivation of Xrn1p. Shown is a northern blot analysis using a probe hybridizing to the exon1 of *NOG2* of RNAs extracted from wild-type (WT), or *xrn1* deletion mutants transformed with plasmids expressing the M1 or M4 constructs (Figure 4A).

In the *rrp6*Δ mutant, the levels of hmsnoRNAs expressed from the M1 or M2 plasmids decreased with an accumulation of unspliced precursors, consistent with our previous conclusion that the nuclear exosome processes the 3′-end of hmsnoRNAs. The *dom34*Δ and *upf1*Δmutations had no major effect on the accumulation of hmsnoRNAs expressed from these plasmids, in contrast to the result described above which showed a decreased accumulation of hmsnoRNAs in the *lea1*Δ*upf1*Δ mutant (Figure 3D). Since inactivating Upf1p does not impact the level of hmsnoRNAs generated by splicing signal mutations, we interpret the decreased accumulation of hmsnoRNA previously observed in the *lea1*Δ*upf1*Δ mutant as the result of indirect effects. This interpretation is supported by previous work showing that mutations that reduce splicing efficiency and inactivate NMD result in synthetic growth defects (33), which may indirectly impair processes required for the accumulation of hmsnoRNAs.

In contrast to all other RNA degradation mutants analyzed previously, a large increase of all *NOG2* RNA species was detected in the *xrn1*Δ mutant deficient for the major cytoplasmic 5′-3′ exonuclease (34)(Figure 4B). While the increased accumulation of spliced and unspliced species was expected, the increase in abundance of the hmsnoRNA strongly suggests that these species are degraded in the cytoplasm by the general mRNA decay pathway, which relies primarily on Xrn1p. As hmsnoRNAs contain the 5′-ends and exon1 sequences of the *NOG2* gene, they may contain a ^7^meG-cap similar to that of mRNAs which would subject them to the general decay pathway for cytoplasmic mRNAs (35). In order to show that hmsnoRNAs are degraded by the general cytoplasmic decay pathway which requires decapping, we analyzed hmsnoRNA accumulation resulting from the M1 mutation in a strain lacking the decapping enzyme Dcp2p. As observed previously for the *xrn1*Δ mutant, inactivation of Dcp2p resulted in an increased accumulation of the hmsnoRNAs (Figure S7). This result is consistent with the idea that hmsnoRNAs are degraded by the major cytoplasmic decay pathway, which involves Dcp1/2-mediated decapping and 5′-3′ degradation by Xrn1p.

### HmsnoRNAs are affected by delocalization of Rat1p in the cytoplasm

To test if hmsnoRNAs are affected by the nuclear 5′-3′ exonuclease Rat1p (also known as Xrn2p), we inactivated Rat1p by anchor away and assessed the accumulation of hmsnoRNAs from the M1 construct (Figure 4C). This analysis revealed higher hmsnoRNAs levels in conditions where Rat1p is depleted from the nucleus, suggesting that a fraction of hmsnoRNAs is nuclear. It is also possible that the inactivation of Rat1p may limit degradation of unspliced precursors as shown previously (25, 28), which could then be converted into hmsnoRNAs and result in higher hmsnoRNA levels.

To assess whether Rat1p could degrade hmsnoRNAs in the cytoplasm, we used plasmids expression wild-type Rat1p, or a mutant lacking its nuclear localization signal (*NLSΔ*) which was shown previously to be able to rescue Xrn1p activity (36). The expression of these constructs had little effect in a wild-type context (Figure 4D). By contrast, expressing the *rat1-NLSΔ* mutant in a strain lacking Xrn1p resulted in a strong decrease of hmsnoRNA, unspliced and mature mRNAs levels, while expressing a wild-type version of Rat1p had little impact. This result shows that a version of Rat1p which is delocalized in the cytoplasm can destabilize hmsnoRNAs in the absence of Xrn1p, providing further evidence that hmsnoRNAs are primarily degraded through cytoplasmic 5′-3′ decay.

### HmsnoRNAs share some of the hallmarks of mature snoRNAs

The plasmid expressing *NOG2* with a mutated 5′-SS provided us with a suitable system to investigate if hmsnoRNAs share some of the features of mature snoRNPs. We first asked if hmsnoRNAs are bound by a snoRNP protein found in mature snoRNPs. We performed immunoprecipitation of RNAs in a strain expressing the M1 mutant and a ZZ-tagged version of Nop10 (37), a core component of H/ACA snoRNPs. A control strain was included which did not express tagged Nop10p. Northern blot analysis of RNAs purified on IgG beads (which bind the ZZ-tagged Nop10p) showed that hmsnoRNAs co-purified with Nop10p, as did the mature forms of *snR191* (Figure 4E, IP lane 2). By contrast, non H/ACA RNAs such as *snR54* or *scR1* ncRNA were not efficiently purified (Figure 4E and Figure S8) and no RNAs were retained on IgG beads in the strain that did not express tagged Nop10p (IP lane 1), showing the specificity of this immunoprecipitation procedure. The ratio of signals for the RNAs found in the Nop10p immunoprecipitates compared to the input showed very similar values for the mature snR191 and the hmsnoRNA (Figure S8), suggesting that hmsnoRNAs are quantitatively bound by Nop10p. Interestingly, unspliced precursors were also immunoprecipitated by Nop10p (Figure 4E). This is consistent with prior work showing co-transcriptional assembly of H/ACA snoRNP proteins on an intron-encoded H/ACA snoRNA in *S*.*cerevisiae* (38).

If hmsnoRNA are bound by snoRNP proteins and stabilized by their binding, one would expect that mutation of snoRNA sequence elements that promote snoRNP assembly would result in a decrease of a steady-state levels of hmsnoRNAs, as described previously for mature H/ACA snoRNAs (1). To test this hypothesis, we generated a variant of the M1 mutant in which the ACA box of *snR191* is deleted (M4 mutant; Figure 4A). This mutation resulted in the complete destabilization of the hmsnoRNA species, which were no longer detectable (Fig.4C, 4F). Strikingly, anchoring away Rat1p from the nucleus partially restored the accumulation of hmsnoRNAs expressed from this construct (Fig.4C). This result shows that defective assembly of snoRNP proteins due to the ACA box deletion triggers degradation of the hmsnoRNAs by nuclear 5′-3′ decay. By contrast, deletion of Xrn1p did not result in any rescue of hmsnoRNA levels from the M4 mutant, as opposed to what was observed for the hmsnoRNAs generated from the M1 construct (Fig.4F). These results show that hmsnoRNAs that are not properly assembled into snoRNPs are primarily degraded by Rat1p in the nucleus and are unaffected by cytoplasmic turnover.

## Discussion

In this work, we show that inactivation of splicing by impairing splicing factors or mutating splicing signals leads to the production of hybrid mRNA-snoRNA species for both box C/D or H/ACA snoRNAs. Extensive mapping of these species by northern blot and Oxford Nanopore long-read sequencing showed that hmsnoRNA have extended 5′-ends containing mRNAs sequences that include 5′-UTR and exon1 and that their 3′-ends match those of mature snoRNAs. This chimeric architecture is reminiscent of that of sno-lncRNAs(17), but hmsnoRNA contain mRNA-like features instead of a snoRNA at their 5′-ends. The data presented above suggest a general pathway for the production and degradation of hmsnoRNAs (Figure S9). Splicing inactivation results in accumulation of unspliced pre-mRNAs, which are bound by at least a subpopulation of snoRNP proteins (eg Nop10p for the *NOG2-snR191* hmsnoRNA) and then trimmed to or near the mature 3′-end of the snoRNA sequence by the nuclear exosome (Figure S9). Early binding of snoRNP proteins is consistent with prior studies that have shown co-transcriptional assembly of snoRNP components (38–41). This assembly can occur in a splicing-independent manner (41), which explains why Nop10p can bind to hmsnoRNAs produced when splicing is inactivated (Figure 4E) and why deletion of the ACA box of a *NOG2-snR191* gene containing a 5′-splice site mutant significantly destabilizes the RNAs generated from this construct and triggers nuclear decay by Rat1p (Figure 4C). After nuclear 3′-end processing by the exosome, hmsnoRNAs are exported to the cytoplasm where they can be degraded by the general 5′-3′ decay pathway that involves Dcp2p and Xrn1p (Figure S9). Further evidence for cytoplasmic localization of hmsnoRNAs is provided by the fact a version of Rat1p mis-localized in the cytoplasm can reduce hmsnoRNA levels in a strain lacking Xrn1p (Figure 4D). Despite degradation by these 5′-3′ decay pathways, hmsnoRNA can accumulate to relatively high levels, from 10 to 64% relative to the mature snoRNA levels, depending on the snoRNA (Table S1).

We found that inactivation of splicing factors involved at different steps of the spliceosome cycle results in the accumulation of hmsnoRNAs. This observation might be unexpected for factors involved in the 2^nd^ catalytic step such as Slu7p or Prp18p, as anchoring away these factors is expected to produce mostly lariat intermediates and cleaved 5′exons, but not unspliced precursors which we showed are converted into hmsnoRNAs by the nuclear exosome. However, a recent study offers an explanation to this conundrum, as inactivation of late splicing factors can result in unspliced precursors accumulation *in vivo* (42), possibly because of recycling defects. Our work sheds light onto previous results that described the accumulation of RNAs similar to hmsnoRNAs upon mutation of the branchpoint sequence of the host intron of *snR18* (previously called U18)(43). The result reported in this previous study is reminiscent of the effect detected when mutating the 5′-SS and the branchpoint of *NOG2*. In the previous study, the 5′-extended *snR18* species were interpreted as intermediates in the processing pathway(43). However, there was no evidence provided that these species could be converted into functional mature products, and based on our observations, it is more likely that they correspond to dead-end products similar to hmsnoRNAs, and which are defective byproducts of splicing inactivation.

The accumulation of RNAs similar to hmsnoRNAs has also been reported when the pre-tRNA processing enzyme RNase P is inactivated by a thermosensitive (ts) mutation (15). These species were considered intermediates in the pathway, and their accumulation was interpreted as evidence for a direct involvement of RNase P in the processing pathway of intron-encoded snoRNAs. However, RNase P-mediated cleavage intermediates could not be detected *in vivo* in this study (15); furthermore, lariat introns containing snoRNAs accumulate at very high levels in a debranching enzyme mutant(10), which argues against a major cleavage pathway of intronic sequences by RNase P. We propose instead that the accumulation of hmsnoRNAs-like RNAs in this RNase P ts mutant is in fact due to an indirect inhibition of splicing, as work published later by the same group showed that this mutation results in the accumulation of unspliced pre-mRNA precursors *in vivo* (44) which might be converted into hmsnoRNAs.

Splicing activity has been shown to be inhibited during stress or non-standard growth conditions(45) which suggests that hmsnoRNAs could be naturally produced in such conditions without experimental interference with the splicing process. To assess whether hmsnoRNAs might be produced in growth conditions that reduce splicing efficiency, we analyzed *NOG2* expression in stationary phase, heat shock conditions or treatment with rapamycin (in a wild-type strain that contains a functional TOR pathway, unlike the anchor away strains described above). None of these conditions resulted in the accumulation of hmsnoRNAs (Figure S10). However, the same conditions are also known to generally reduce ribosome biogenesis and snoRNP assembly. Therefore, the absence of detection of these species in these conditions is difficult to interpret, as it might be due to conditions that globally repress ribosome biogenesis and prevent assembly of snoRNP proteins on the hmsnoRNA species.

The results presented here provide a general framework that underscores the importance of the splicing process for the biogenesis of snoRNAs. In the absence of splicing, not only are intron-encoded snoRNAs not processed properly, but the RNAs species that are improperly generated are subject to a degradation pathway similar to that which targets mRNAs in the cytoplasm (Figure S9). A parallel model was proposed to underscore the importance of 5′-end processing for independently transcribed snoRNAs in *S*.*cerevisiae* (18). In the absence of co-transcriptional cleavage of snoRNA precursors by the endonuclease Rnt1p, unprocessed snoRNAs accumulate in the cytoplasm and are not functional(18). Overall, the results described here combined to those reported by Kufel, Proudfoot and colleagues converge to establish a unified model for the importance of RNA processing for the fate of snoRNAs. Regardless of the precise mode of expression and of the nature of the transcription units that produce snoRNAs, RNA processing of snoRNA precursors serves two major purposes: these reactions not only remove flanking sequences, but also dictate the proper fate of snoRNP particles and their mode of degradation. In the case of intron-encoded snoRNAs, the work described here establishes an additional functional role for splicing reactions, beyond simply removing intervening sequences of mRNAs. Finally, these data may shed some light onto the molecular effects of mutations of splicing factors, which can result in a variety of human diseases. While the impact of these mutations on mRNA splicing and alternative splicing patterns has been investigated, their effect on the production of intron-encoded snoRNAs has not been well characterized. It is possible that some of the deleterious effects of these mutations that are causal to disease may be linked to the defective processing of intron-encoded snoRNAs and the production of hmsnoRNA-like species, which may be detrimental to RNA metabolism.

## Materials and Methods

Yeast strain construction and manipulation, RNA extraction and northern blot analysis using riboprobes was performed as described in Wang et al (46). The list of strains, plasmids and oligonucleotides used is provided in Supporting information. Wild-type and knockout yeast strains are from the BY4742 genetic background(47) and knockout strains were obtained from the systematic knockout collection(48). Strains used for the anchor away experiments are from the HHY168 genetic background (20). The detailed protocol used for direct RNA sequencing is described in detail in Supporting information and sequencing data is available at NCBI as BioProject ID: PRJNA827814. For the expression of *NOG2* mutants, the wild-type *NOG2* gene was cloned in pUG35(49) and expressed under the control of its own promoter to create pCL1. The M1, M2 and M4 mutants were created by site-directed mutagenesis from pCL1. For immunoprecipitation of ZZ-tagged Nop10p, we used strains transformed with the pFH35 plasmid that expresses ZZ-tagged Nop10p(37). RNA immunoprecipitation was performed as described in Supporting information.

## Acknowledgements

We thank Anthony Henras for sharing the Nop10-ZZ tagged expression plasmid and for helpful discussions, and Jeff Coller and Arlen Johnson for sharing strains and plasmids. This work was supported by NIGMS grant R35 GM130370 to G.F.C. MRG was supported by NIGMS grant IRACDA 2K12 GM106996-06.

