## Supplemental Methods Figures and Tables for "Splicing inactivation generates hybrid mRNA-snoRNA transcripts targeted by cytoplasmic RNA decay"

**Abstract**

Many small nucleolar RNAs (snoRNA)s are processed from introns of host genes, but the importance of splicing for proper biogenesis and the fate of the snoRNAs is not well understood. Here we show that inactivation of splicing factors or mutation of splicing signals leads to the accumulation of partially processed hybrid mRNA-snoRNA transcripts (hmsnoRNA). HmsnoRNAs are processed to the mature 3′-ends of the snoRNAs by the nuclear exosome and bound by snoRNP proteins. HmsnoRNAs are unaffected by translation-coupled RNA quality control pathways, but they are degraded by the major cytoplasmic exonuclease Xrn1p due to their mRNA-like 5′-extensions. These results show that completion of splicing is required to promote complete and accurate processing of intron-encoded snoRNAs and that splicing defects lead to degradation of hybrid mRNA-snoRNA species by cytoplasmic decay, underscoring the importance of splicing for the biogenesis of intron encoded snoRNAs.

### Supporting Materials and Methods

#### Oxford Nanopore Sequencing

The Saccharomyces cerevisiae Slu7p anchor-away strains was used as a source of RNA. The strain was grown in YPD and when OD600 reached 0.5, Slu7p was depleted from the nucleus by addition of rapamycin to a final concentration of 1ug/ml and incubated for 1 hour. 40 ug of total RNAs extracted from these cells were treated with DNAse I (Invitrogen, catalog #: 18-068-015) according to the manufacturer's protocol. DNase I treated RNAs were incubated with Shrimp Alkaline Phosphatase (NEB, catalog #: M0371S) in order to remove 3' phosphates. This was followed up by treatment with Terminator™ exonuclease (Lucigen, catalog #: MA246E) digestion, as specified in the manufacturer's protocol, to degrade RNAs with 5' monophosphates. The remaining RNAs were then in vitro polyadenylated using E. coli Poly(A) Polymerase (NEB, catalog #: M0276S). Phenol chloroform extraction and ethanol precipitation was performed after each enzymatic treatment.

RNA libraries were prepared from 500 ng of in vitro polyadenylated RNAs using the direct RNA sequencing kit from Oxford Nanopore (ONT, catalog #: SQK-RNA002) as per the manufacturer's instructions. Sequencing was performed using R9.4 flow cells on a MinION Mk1B device and sequenced for 48 hours. Basecalling was performed using Guppy Basecaller (Version 6.1.1+1f6bfa7f8). Reads were then mapped to the Saccharomyces cerevisiae genome (S288C_reference_sequence_R64-3-1) using Minimap 2 (Version 2.17-r941). Reads were visualized using IGV (Version 2.12.3) and figures were prepared using Inkscape (Version 1.1.2).

Data is available as unprocessed FAST5 files, FASTQ files and aligned bam files at NCBI as BioProject ID: PRJNA827814.

#### RNA immunoprecipitation

Yeast cells corresponding to 400 OD600 units of culture were harvested in exponential phase, washed and resuspended in lysis buffer (20mM Tris-HCl pH=8, 300mM K-Acetate, 5mM $MgCl_2$, 1mM Dithiothreitol, 0.2% Triton X-100, protease inhibitors cocktail tablet (Roche) ). Cells were lysed by vortexing in the presence of glass beads (425-600 µm) for 5 minutes. Whole cell lysates were collected after centrifugation at 15,000 rpm for 20 minutes. About 1000µL of lysate and 50µL of pre-washed mouse IgG conjugated magnetic beads (Cell Signaling) were incubated for one hour on a shaker at 4°C. After purification of the beads on a magnetic rack, beads were washed five times with 1000µL of lysis buffer and RNA extraction after the final wash were performed using phenol: chloroform: iso-amyl alcohol solution (VWR).

### Supporting Figures

#### Figure S1. Analysis of hmsnoRNA 3’ ends using 3’ RACE

**(A)** Schematic of 3’ RACE procedure. RNA with an intron-encoded snoRNA is depicted. The RNA was in vitro polyadenylated and an oligo d(T)-containing primer (prMG1) was annealed to the poly-A tail (top panel). RNA was reverse transcribed to cDNA and amplified by PCR (lower panel). Primers prMG3 and prMG5 anneal to the junction between exon 1 and the upstream intron while primers prMG4 and prMG6 anneal to the intron. Primer prMG2 anneals to the 5’ sequence of prMG1, added to the 3’ end of cDNA during reverse transcription.

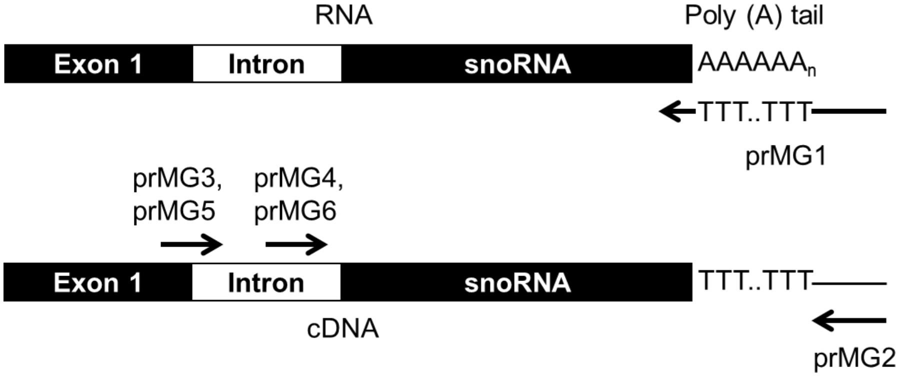


**(B-C)** Sequences of the amplified 3’ ends of hmsnoRNAs from NOG2/snR191 **(B)** and IMD4/snR54 **(C)** obtained from RNAs extracted from a Slu7p-anchor away strain post-rapamycin treatment. Sequences aligning to the snoRNA are bolded, and the poly-A sequence added by in vitro polyadenylation is italicized.

**B.**

TNTTNCNTAN**CCTTTTTGTCAGGGTGCTTCTCTATCCGTTTTAGGATAAACTTATC**
**TACAGAACTGTCTGTTACGGTTTTGGAGGANTAATACTGTTCCCTTATCCCTATT**
**TCCCCGTTCTGGGGAACCCTCATGGGTAAATTANAATGACAACCTTTGTTTTAA**
**AGGTATACCTTCGCTTTTTANAACAGCGAGGATCTTATGAGTTGAGCTTTTGTTA**
**TTTGAGACTTTATCTCGGGCTCCATACAATATGTTCCTACTAAAGATCCTCACAA**
**TT***AAAAAAAAAAAAAAAAAAA*

**C.**

GTGAANGATC**TAAGATGATGATCAACTTTTTATATCAATAACTTTCGTTCTACTGA**
**CTGTGATCAAACGATCTTGTAGAGAACTTTTACTCTGAAT**T*AAAAAAAAAAAAAA*
*AAAAAAA*

**Figure S2**. **Mapping and Polyadenylation status of the *ASC1-snR24* hmsnoRNAs.**

**A.** Structure of the ASC1/snR24 gene and location of the probes used for differential northern blot analysis in panel B. Legends as in Figure 1A. Boxes and line lengths are not to scale.

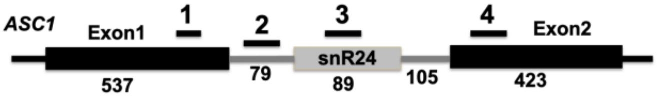


**B**. Northern blot analysis of ASC1/snR24 in the Slu7p-AA strain using probes hybridizing to the indicated regions of the ASC1/snR24 gene. An ethidium bromide staining of the 25S rRNA is shown as a loading control. The *ASC1-snR24* hmsnoRNA species were detected by the 5′-exon and snoRNA probes (probes 1 and 3), and by an intronic probe located 5′ to the snoRNA (probe 2), but not to a probe hybridizing to the 3′-exon (probe 4). As expected, probe 1 also detected the cleaved 5′-exon intermediate, and probes 2-4 detected the lariat intermediate in the Slu7p-AA strain treated with rapamycin. Labeling of the species as in main Figure 1.

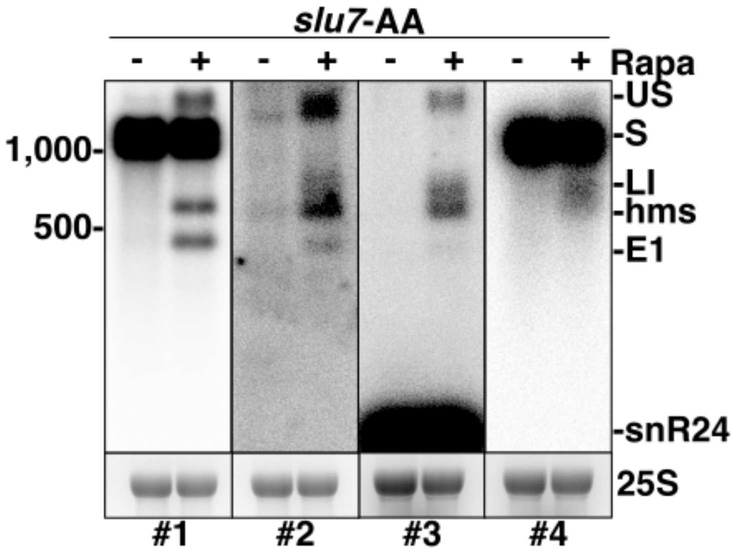


**C.** Analysis of the polyadenylation status of ASC1/snR24. Shown is a northern blot of ASC1/snR24 using a 5′exon probe (#1 in Figure 2C) of RNAs extracted from the Slu7p-FRB tagged strain grown after treatment with rapamycin. T = total RNAs; pA+= polyadenylated RNAs selected by oligodT affinity; pA- = Non-polyadenylated RNAs extracted from the supernatant of the oligodT affinity purification. scR1 was used as a non-polyadenylated RNA negative control.

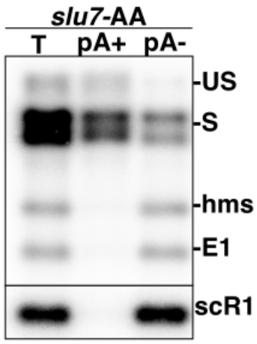

**Figure S3**. Oxford Nanopore sequencing reads obtained from the slu7-anchor away strain for the RPL7B and RPS22B regions. Labeling of the species as in main Figure 2.

**A. RPL7B Region.**

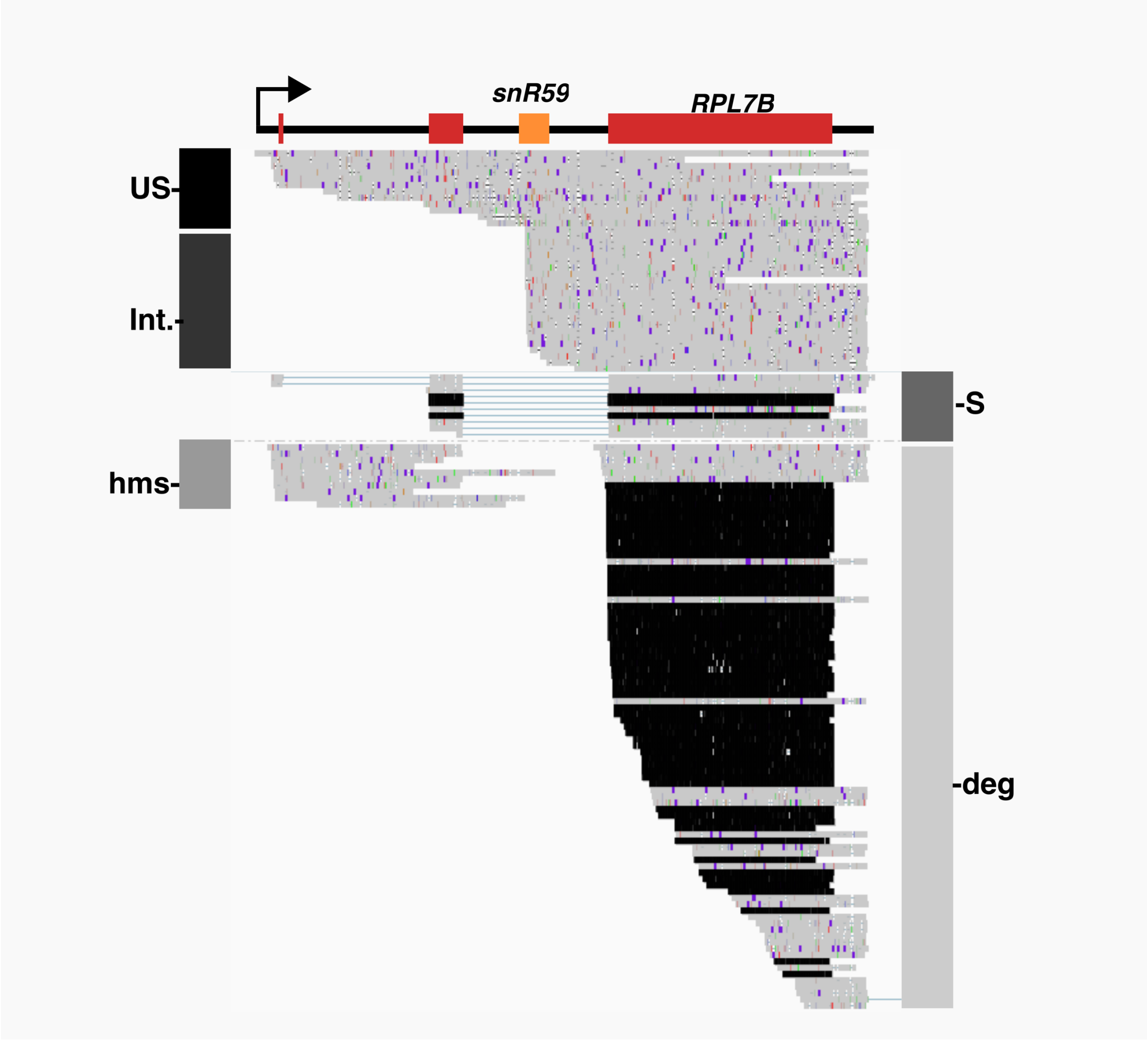


**B. RPS22B Region.**

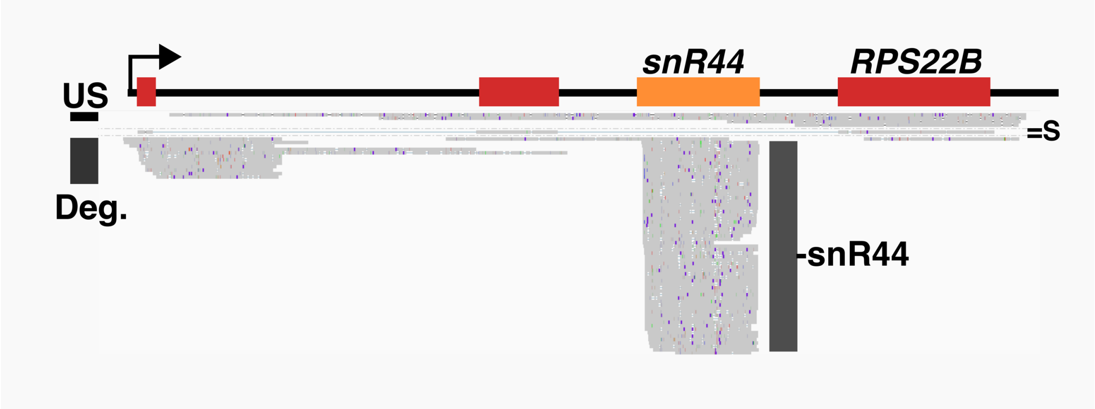

**Figure S4**.
Northern blot analysis of *NOG2/snR191* using a probe complementary to the exon1 of *NOG2* or a probe complementary to snR191 in strains expressing anchor-away versions of Rrp6p, Slu7p, or both Slu7p and Rrp6p. Strains in the name of the gene italicized indicate that the corresponding protein was FRB-tagged (eg *rrp6* = *rrp6-FRB*). Each FRB tagged strain was grown in normal medium (-Rapa) or shifted for 1hr in a medium containing Rapamycin (+Rapa) to promote export of the corresponding proteins out of the nucleus. GAPDH was used as a loading control. Labeling of the species as in Main Figure 1. scR1 was used as a loading control.

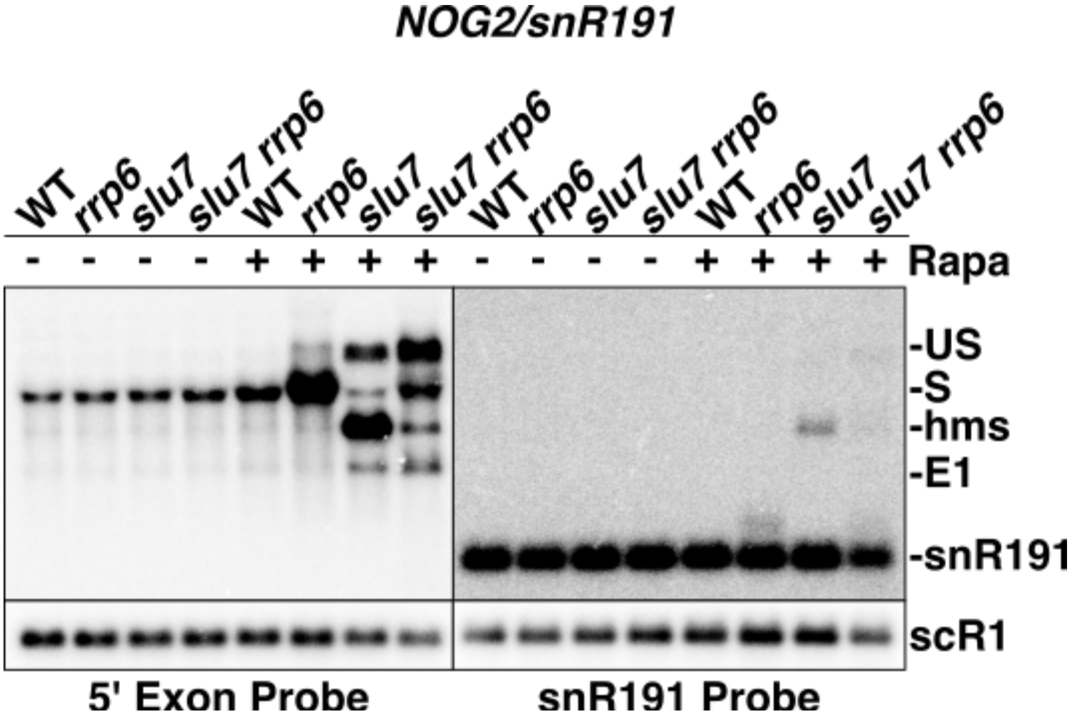

**Figure S5**. Northern blot analysis of *NOG2* in a wild-type strain transformed with a centromeric vector (pUG35) or pUG35 containing a *NOG2* gene insert.

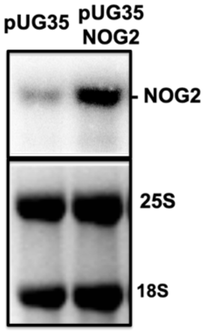


The *NOG2* gene inserted in the pUG35 plasmid is expressed under the control of its endogenous promoter. The top panel shows a northern blot analysis of NOG2 using a probe hybridizing to the exon1 of NOG2. The bottom panel shows an Ethidium bromide staining of rRNAs from the gel used for the northern blot shown above. The *NOG2* label indicates the spliced *NOG2* mRNA.

**Figure S6**. Northern blot analysis of *NOG2* M1 and M2 mutant expression in wild-type and mutant strains grown at 20°C or 30°C. Wild-type (WT) or the indicated deletion mutants were transformed with the pUG35 vector or the pUG35 plasmids expressing mutants M1 or M2 (Main Figure 4), grown at 20°C or 30°C, and analyzed by northern blot using a probe hybridizing to the exon1 of NOG2. The different RNA species are labeled as in Fig.S2.

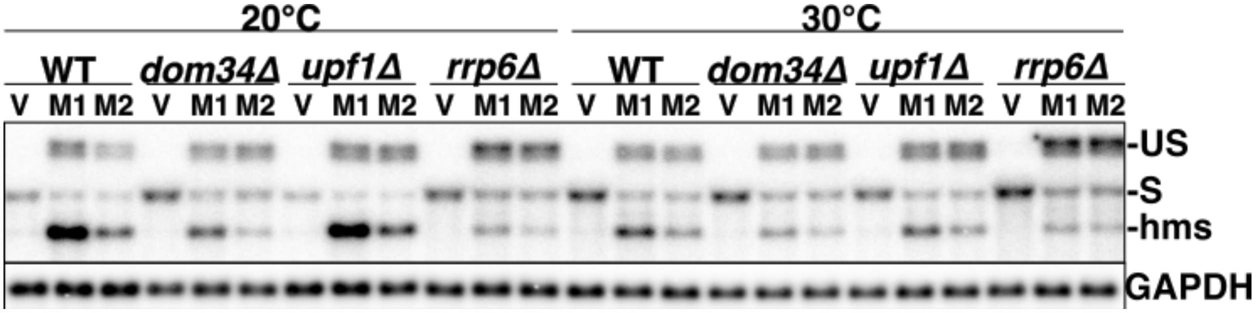


**Figure S7.**

Northern blot analysis of *NOG2* M1 mutant expression in wild-type, *xrn1*Δ and *dcp2*Δ mutant strains. The probe used hybridized the exon1 of NOG2. An ethidium bromide staining of the 25S rRNA was used as a loading control. The different RNA species are labeled as in Fig S2.

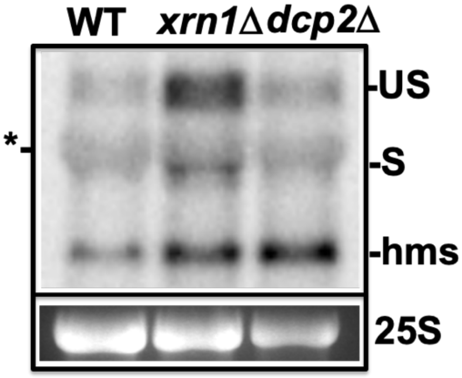

**Figure S8**. Quantifications of the ratios of RNA species detected in the Nop10 immunoprecipitations shown in Figure 4C vs. the RNAs detected in the input fractions.

Ratios are shown for the two independent biological replicates. The scR1 RNA was not analyzed for Replicate 1.

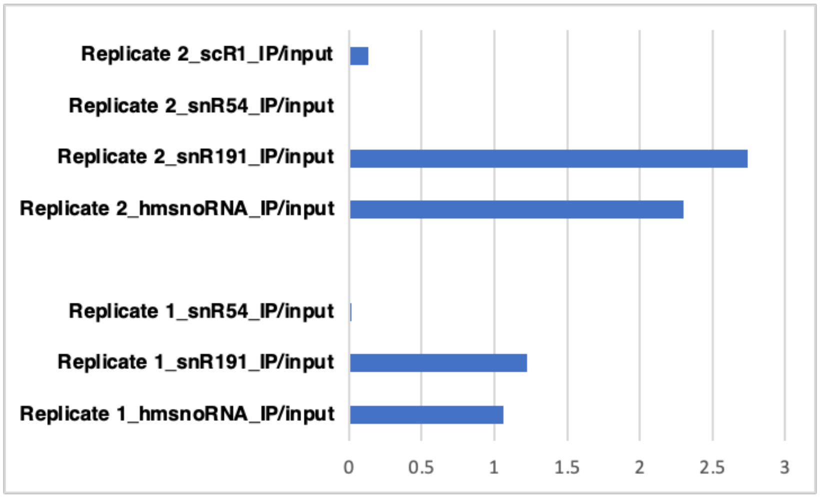


**Figure S9.** Model of biogenesis of intron-encoded snoRNAs and impact of splicing inhibition on intron-encoded snoRNAs processing and degradation.

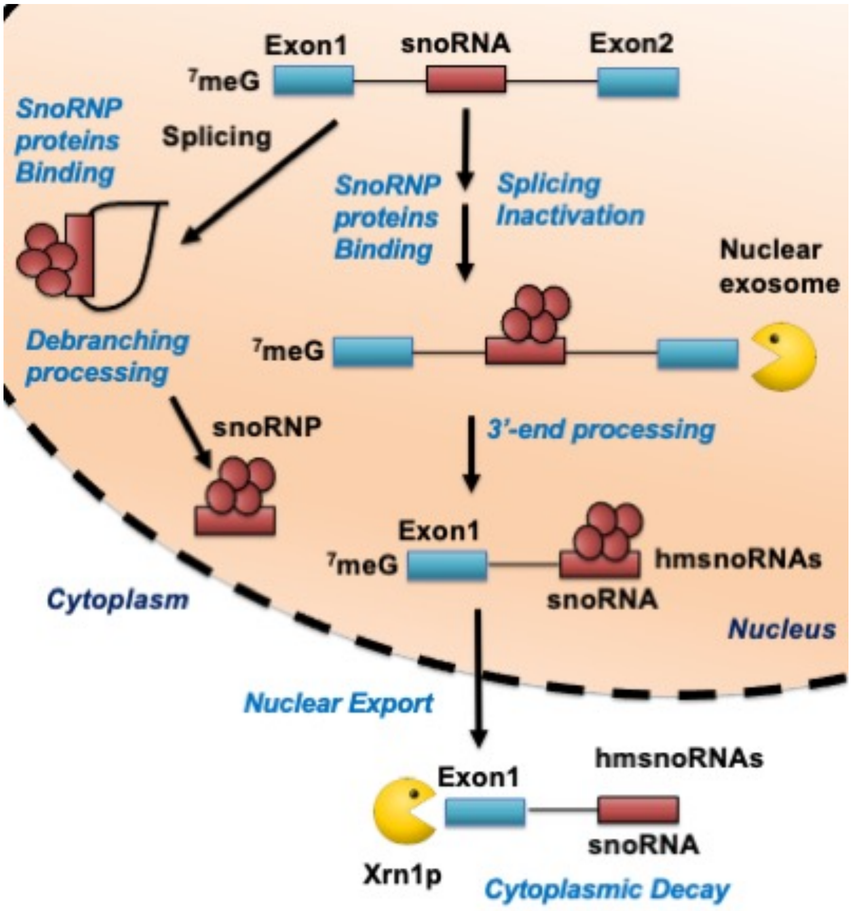

**Figure S10. Analysis of NOG2 expression in non-standard growth conditions.**

Shown are northern blots of *NOG2* of RNAs extracted from wild-type yeast strain grown in log phase (log), stationary phase (stat), exposed for 2hours of heat shock (39°C HS) or treated for one hour with Rapamycin. None of the conditions used resulted in the production of hmsnoRNAs. Only the spliced form of *NOG2* (S) is indicated.

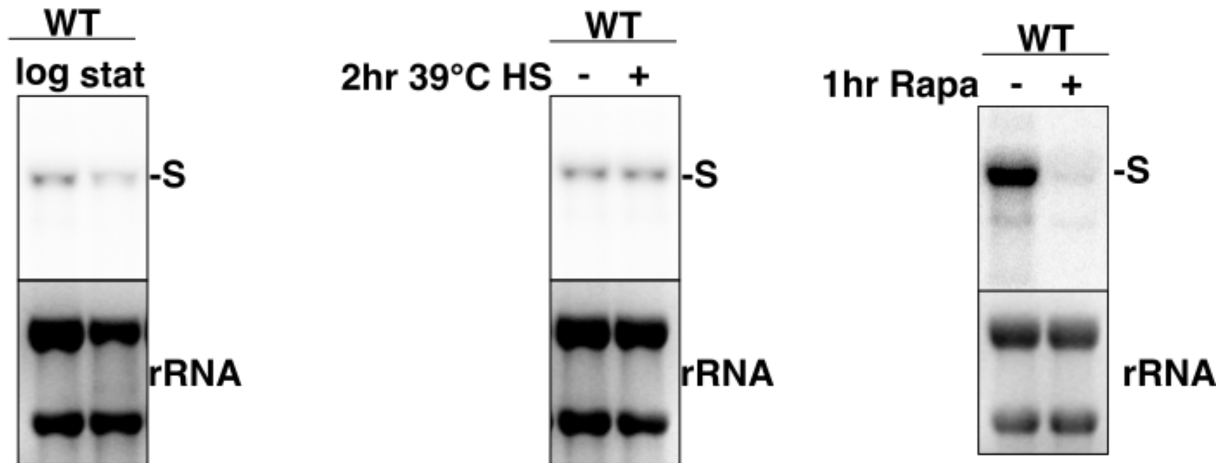

**Table S1: Quantification of the abundance of hmsnoRNAs relative to the mature snoRNAs based on northern blots using snoRNA probes.**

| Gene | snoRNA | HmsnoRNA/snoRNA Ratio | Method | Replicates |
|---|---|---|---|---|
| NOG2 | snR191 | 19.2% | Slu7-Anchor Away | 1 |
| NOG2 | snR191 | 11.5% | Slu7-Anchor Away | 2 |
| NOG2 | snR191 | 16.0% | *lea1Δ* | 1 |
| NOG2 | snR191 | 16.4% | *lea1Δ* | 2 |
| ASC1 | snR24 | 15.4% | Slu7-Anchor Away | 1 |
| IMD4 | snR54 | 69% | Slu7-Anchor Away | 1 |
| TEF4 | snR38 | 34.7% | Slu7-Anchor Away | 1 |

**Table S2: Strains used in this study**

| Strain ID | Genotype | Source |
|---|---|---|
| yCL1 (BY4742) | MATα his3Δ1 leu2Δ0 lys2Δ0 ura3Δ0 | Ref. 47 |
| yCL2 | CL1 with *lea1Δ::KANR* | Ref. 48 |
| yCL3 | CL1 with *isy1Δ::KANR* | Ref. 48 |
| yCL4 | CL1 with *lea1Δ::KANR dom34Δ::HIS* | This study |
| yCL5 | CL1 with *lea1Δ::KANR upf1Δ::HIS3* | This study |
| yCL6 | CL1 with *lea1Δ::KANR rrp6Δ::HYGR* | This study |
| yCL7 | CL1 with *dom34Δ::KANR* | Ref. 48 |
| yCL8 | CL1 with *upf1Δ::KANR* | Ref. 48 |
| yCL9 | CL1 with *rrp6Δ::KANR* | Ref. 48 |
| yCL10 | CL1 with *xrn1Δ::KANR* | Ref. 48 |
| yCL11 (HHY168) | MATa ade2-1 can1-100 his3-11,15 leu2-3,112 trp1-1 ura3-1 tor1-1 *fpr1Δ::NAT* RPL13A-2xFKBP12::TRP1 | Ref.20 |
| yCL12 | CL11 *SLU7-FRB::KanMX6* | This study |
| yCL13 | CL11 *PRP16-FRB::KanMX6* | This study |
| yCL14 | CL11 *PRP18-FRB::KanMX6* | This study |
| yCL15 | CL11 *PRP22-FRB::KanMX6* | This study |
| yCL16 | CL11 *PRP5-FRB::KanMX6* | This study |
| yCL17 | CL11 *PRP28-FRB::KanMX6* | This study |
| yCL18 | CL11 *RRP6-FRB::His PRP18-FRB::KanMX6* | This study |
| yCL19 | CL11 *RRP6-FRB::His SLU7-FRB::KanMX6* | This study |

**Table S3: Plasmids used in this study**

| Plasmid ID | Vector |
|---|---|
| pCL1 | pUG35 |
| pCL2 | pCL1 with NOG2 |
| pCL3 | pCL1 with *nog2-mutated 5'splice site* |
| pCL4 | pCL1 with *nog2-mutated 5'splice site and branchpoint* |
| pCL5 | *pCL1 with nog2-mutated 5'splice site and ACA deletion* |
| pAJ203 | *CEN LEU2 RAT1* (WT) (Ref.36) |
| pAJ228 | *CEN LEU2 rat1-NLSΔ* (Ref.36) |
| pFH35 | *CEN LEU2 NOP10-ZZ (Ref.37)* |

**Table S4: Oligonucleotides List.**

| Name | Sequence (5'→3') |
|---|---|
| NOG2_FWD | CGTTGGTTCGGTAACACAAG |
| *Usage* | *For synthesis of riboprobes binding to NOG2 exon 1* |
| NOG2_REV_T3 | AATTAACCCTCACTAAAGGGAGTAAATGCTTGTGTGGTGTTTCC |
| *Usage* | *For synthesis of riboprobes binding to NOG2 exon 1* |
| snR191_FWD | CAAACCTTTTTGTCAGGGTGC |
| *Usage* | *For synthesis of riboprobes binding to snR191* |
| snR191_REV_T3 | AATTAACCCTCACTAAAGAATTGTGAGGATCTTTACTACGAAC |
| *Usage* | *For synthesis of riboprobes binding to snR191* |
| NOG2_intron_upsnR_oligo | TCTCTCGTGCTATCCTCTTGGTTGGAAAGAATC |
| *Usage* | *For synthesis of oligoprobes binding to the intronic region upstreamsnR191* |
| NOG2_Exon2_FWD | CATTTGTCAAAGGAACGTCC |
| *Usage* | *For synthesis of riboprobes binding to NOG2 exon 2* |
| NOG2_Exon2_REV_T3 | AATTAACCCTCACTAAAGTACCTCTATGCCGTCTTCTC |
| *Usage* | *For synthesis of riboprobes binding to NOG2 exon 2* |
| IMD4_FWD | TTCAAATCATGCTTGGCTCATC |
| *Usage* | *For synthesis of riboprobes binding to IMD4 exon 1* |
| IMD4_REV_T3 | AATTAACCCTCACTAAAGACTGGGAAGCCAGAGAAACC |
| *Usage* | *For synthesis of riboprobes binding to IMD4 exon 1* |
| TEF4_FWD | TTTGGCCCTCGATAGATTCA |
| *Usage* | *For synthesis of riboprobes binding to TEF4 exon 1* |
| TEF4_REV_T3 | AATTAACCCTCACTAAAGGGAGTGGATAGCCAAAGCTTCAGT |
| *Usage* | *For synthesis of riboprobes binding to TEF4 exon 1* |
| ASC1_FWD | ATGGCATCTAACGAAGTTTTAGTT |
| *Usage* | *For synthesis of riboproes binding to ASC1 exon 1* |
| ASC1_REV_T3 | AATTAACCCTCACTAAAGGGAGCTTAACCATTTTGTCGTTACCGG |
| *Usage* | *For synthesis of riboproes binding to ASC1 exon 1* |

| | |
|---|---|
| snR54-IMD4 F | CCC AGT TAC TGG TAT GTT AT |
| *Usage* | *For synthesis of riboprobes binding to snR54* |
| snR54-IMD4 T3 R | AAT TAA CCC TCA CTA AAG GGA CGT TTG ATC ACA GTC AGT AG |
| *Usage* | *For synthesis of riboprobes binding to snR54* |
| snR38-TEF4 F | GGCTATCCAATTTTATTGTA |
| *Usage* | *For synthesis of riboprobes binding to snR38* |
| snR38-TEF4 T3 R | AATTAACCCTCACTAAAGGGGGTTACCTATTATTACCCAT |
| *Usage* | *For synthesis of riboprobes binding to snR38* |
| NOG2_SalI_F | CGC GCG GTC GAC GCAGGCCAACAAACTGAACG |
| *Usage* | *For creation of pCL2* |
| NOG2_SacI_R_pUG35 | CGC GCG GAG CTC AGCCCTGATAACGAGACACG |
| *Usage* | *For creation of pCL2* |
| NOG2_5'ss_mutation_F | GGTACCTACCTGGGTTGCAAGCCACTAACTTTTTTTTTTTTTTTTTTGATTCTTTCC |
| *Usage* | *For creation of pCL3* |
| NOG2_5'ss_mutation_R | GGAAAGAATCAAAAAAAAAAAAAAAAAAGTTAGTGGCTTGCAACCCAGGTAGG |
| *Usage* | *For creation of pCL3* |
| NOG2_bp_mutation_F | GGCAAAAGATTACCTATTATGAAACAAAATGTATTTTTCCTGCCTAGAAGGTTTTTTTTT |
| *Usage* | *For creation of pCL4* |
| NOG2_bp_mutation_R | CCTACTTGTAAAGTTAGAATTGAAAAAAAAAAACCTTCTAGGCAGGAAAAATACATTTTG |
| *Usage* | *For creation of pCL4* |
| snR191_ACA_del_F | CCA TAC AAT ATG TTC GTA GTA AAG ATC CTC ATT TTC TAC TTT TTT TTT TTT TTT AGT TTA |
| *Usage* | For creation of pCL5 |
| snR191_ACA_del_R | TAA ACT AAA AAA AAA AAA AAA GTA GAA AAT GAG GAT CTT TAC TAC GAA CAT ATT GTA TGG |
| *Usage* | For creation of pCL5 |
